# Deep neural network analysis employing diffusion basis spectrum imaging metrics as classifiers improves prostate cancer detection and grading

**DOI:** 10.1101/2021.03.22.436514

**Authors:** Zezhong Ye, Qingsong Yang, Joshua Lin, Peng Sun, Chengwei Shao, Yongwei Yu, Luguang Chen, Yasheng Zhu, Anthony T. Wu, Ajit George, Chunyu Song, Ruimeng Yang, Jie Zhan, Sam E. Gary, Jeffrey D. Viox, Zhen Wang, Minjie Wang, Yukun Chen, Eric H. Kim, Joseph E. Ippolito, Jianping Lu, Sheng-Kwei Song

**Author notes:** These authors contributed equally to this work (Zezhong Ye, Qingsong Yang). **Correspondence address to:** Sheng-Kwei Song, Ph.D., Department of Radiology, Washington University School of Medicine, Room 3221, 4525 Scott Ave. St. Louis, MO 63110, USA., E-mail address, Jianping Lu, M.D., Department of Radiology, Changhai Hospital, Shanghai, China.

## Abstract

Structural and cellular complexity of prostatic histopathology limits the accuracy of noninvasive detection and grading of prostate cancer (PCa). We addressed this limitation by employing a novel diffusion basis spectrum imaging (DBSI) to derive structurally-specific diffusion fingerprints reflecting various underlying prostatic structural and cellular components. We further developed diffusion histology imaging (DHI) by combining DBSI-derived structural fingerprints with a deep neural network (DNN) algorithm to more accurately classify different histopathological features and predict tumor grade in PCa. We examined 243 patients suspected with PCa using *in vivo* DBSI. The *in vivo* DBSI-derived diffusion metrics detected coexisting prostatic pathologies distinguishing inflammation, PCa, and benign prostatic hyperplasia. DHI distinguished PCa from benign peripheral and transition zone tissues with over 95% sensitivity and specificity. DHI also demonstrated over 90% sensitivity and specificity for Gleason score noninvasively. We present DHI as a novel diagnostic tool capable of noninvasive detection and grading of PCa.

**One sentence summary:** Diffusion histology imaging noninvasively and accurately detects and
grades prostate cancer.

## Introduction

Prostate cancer (PCa) is the most common and the second most deadly cancer among men in the United States (*1*). Over the last 30 years, PCa screening of asymptomatic men using serum prostate-specific antigen (PSA) blood testing has remained the standard of care for early detection for PCa (*2*). Men with elevated serum PSA are sent for transrectal ultrasound (TRUS)-guided biopsy for diagnosis and grading of PCa (*3*). Considered a generally safe and well-tolerated outpatient procedure (*4*), needle biopsy is potentially associated with complications affecting patients’ quality of life to various degrees (*5, 6*). Unfortunately, over 50% of biopsies performed on men with elevated PSA were cancer free (*7*), and 30-50% of the detected cancers were low grade (*8*). A negative biopsy does not exclude the presence of PCa since biopsy can miss tumor foci. subsequent transrectal prostate biopsy increases the risk of infectious complications (*9*), which is a major barrier for patients to remain on active surveillance (*10*).

To address these limitations, prostate multiparametric magnetic resonance imaging (mpMRI) has been increasingly used to risk stratify men prior to prostate biopsy (*11, 12*). Prostate mpMRI allows for the visualization of lesions that can be identified and assessed for the likelihood of clinically-significant PCa using the Prostate Imaging-Reporting and Data System (PI-RADS) (*13, 14*). Additionally, a prostate biopsy specifically targeting these MRI-defined “tumors” can be performed (e.g. MRI-targeted biopsy). Thus, prostate MRI is a particularly promising tool for active surveillance to minimize the frequency of repeat biopsies (*15, 16*).

Owing to the lack of pathological specificity of mpMRI-defined lesions, inflammation and BPH can confound the interpretation of prostate MRI resulting in high variability across individual radiologists (*17-20*). In fact, biopsy of mpMRI-defined prostate lesions are benign in over 35% of cases (i.e. false positive rate), and biopsy of normal mpMRI areas are found to harbor aggressive PCa in over 15% of patients (i.e. false negative rate) (*11, 21*). Thus, prostate mpMRI cannot eliminate unnecessary initial biopsies in men with suspected PCa (e.g. elevated PSA), and cannot reduce the frequency of repeat biopsies for men on active surveillance (*22, 23*).

The greatest challenge to accurate prostate MRI interpretation remains the lack of specificity of MRI signals to identify PCa, due to other prostate tissue signals that can “mimic” PCa according to PI-RADS criteria (e.g. inflammation) (*17, 18*). We previously developed diffusion basis spectrum imaging (DBSI), which utilizes a data-driven multiple-tensor modeling approach to deconvolute cellular and structural profiles within an image voxel (*24*). DBSI-derived structural metrics distinguish and quantify various pathologies in an array of central nervous system (CNS) disorders (*25-29*). In this study, we examine whether DBSI-derived metrics reflect structural and cellular changes associated with PCa. To improve the PCa diagnostic accuracy, we have further developed a novel diffusion histology imaging (DHI) approach (*28*) using artificial intelligence algorithms (*30, 31*) to recognize underlying structural and pathological signature patterns of cancer based on DBSI-derived structural metrics, an equivalent of filtering nuisance confounds in raw data, to predict the PCa Gleason Grade Group (GG). In this proof-of-concept study, we will test the feasibility of performing deep neural network analysis on DBSI structural metrics derived from diffusion-weighted MRI data for detecting and grading PCa.

## Results

### Patients

Of the 243 patients included in this study (Table S1 and Figure S1), 93 had a PIRADS score ≤ 3; 54 were biopsy-negative of PCa; and 96 were biopsy positive of PCa.

### DBSI derived diffusion metrics distinguish tissue structures in prostatectomy specimens

A prostatectomy specimen from one PCa patient was imaged (Fig. 1a). T2-weighted images and the apparent diffusion coefficient (ADC) maps from conventional prostate MRI did not accurately reflect structural heterogeneity seen in H&E images (Fig. 1a; i, ii, iii, iv). Coexisting histopathological components in the specimen exhibited distinct DBSI-metric profiles: lymphocytes (Fig. 1a; iii, iv, lower left squares) associate with highly-restricted diffusion, i.e., DBSI-derived isotropic ADC < 0.1 µm^2^/ms (Fig. 1b); epithelium and tumor cells (Fig. 1a; i, iv, lower right squares) associated with restricted diffusion (Fig. 1b; 0.1 ≤ DBSI-isotropic ADC < 0.8 µm^2^/ms), stroma (Fig. 1a; ii) associated with hindered diffusion (Fig. 1b; 0.8 ≤ DBSI-isotropic ADC < 1.5 µm^2^/ms), and intra-luminal space (Fig. 1a; i, iii) associated with free diffusion (Fig. 1b; 1.5 ≤ DBSI-isotropic ADC < 2.0 µm^2^/ms). DBSI metric profiles of each image voxel reflect the coexistence of heterogeneous histological components appearing as multiple signature peaks (Fig. 1b). Collagen fibers were modeled by an anisotropic diffusion tensor while lymphocytes, epithelial cells, stroma (without coherent orientation) and luminal contents were each reflected by highly-restricted, restricted, hindered and free isotropic diffusion tensors, respectively (Fig. 1c). Heat maps of DBSI-derived diffusion tensor fractions of the specimen can thus highlight the spatial distribution of various cellular and structural components of PCa as well as adjacent prostate tissue (Fig. 1d).

**Figure 1.**
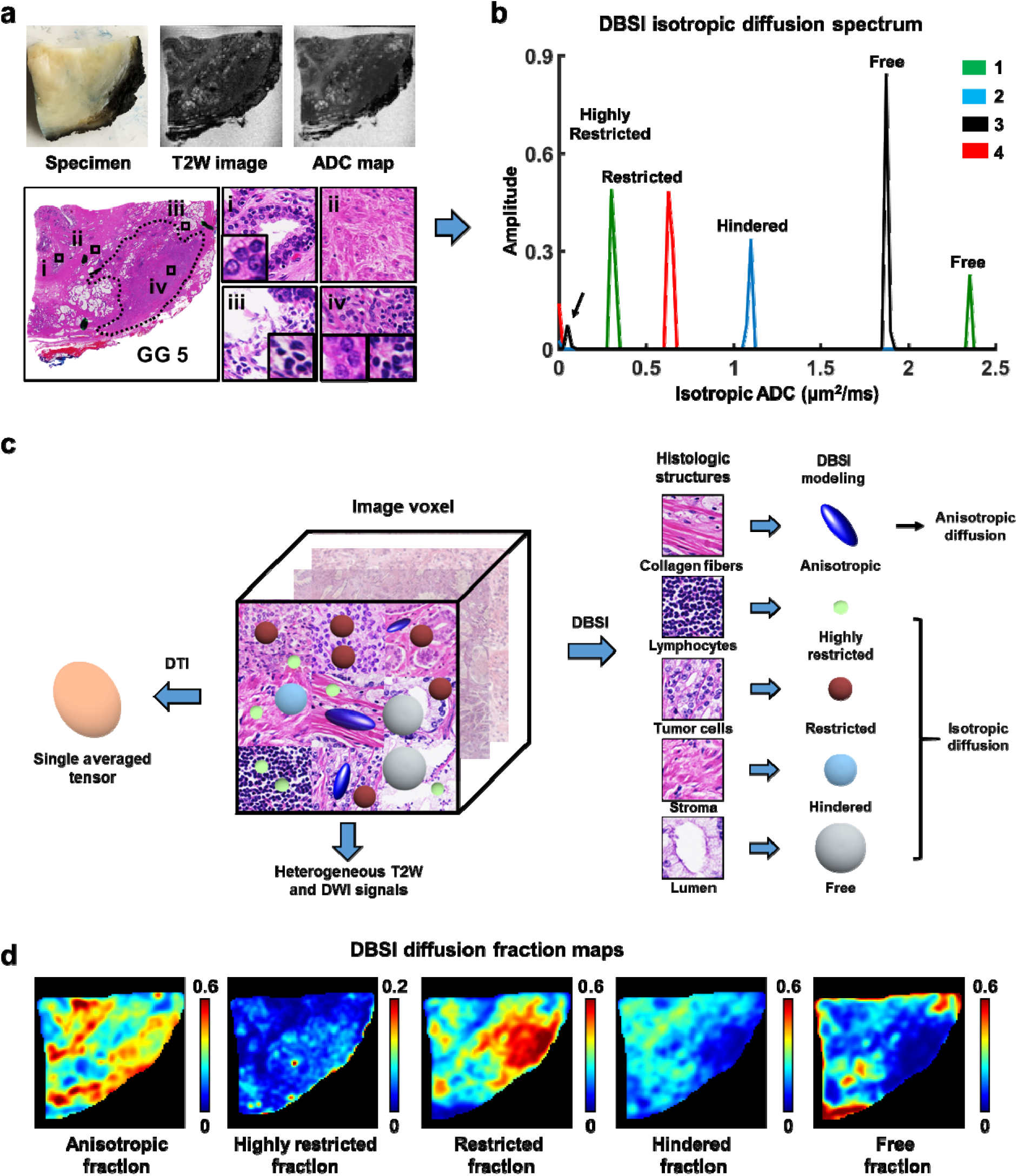
DBSI detected different histopathological structures based on their diffusion signatures. (a) The *ex vivo* MR images and the corresponding H&E images from one representative step-sectioned prostate specimen were shown. ROIs containing stroma, inflammation and PCa were highlighted as follows: lymphocytes (a, iii &iv, lower right square), PCa cells (a, I &iv, lower left square) and stroma tissue (a, ii) were associated with respective signatures: highly-restricted (0 - 0.1 m^2^/ms, lymphocytes), restricted (0.1 - 0.8 m^2^/ms, PCa) and hindered (0.8 - 1.5 m^2^/ms, *stroma*) diffusion. (b) Distinct diffusion patterns, or “structural fingerprints”, from various histopathological structures were identified in each of the ROIs (ROI i: green peaks; ROI ii: blue peaks; ROI iii: black peaks; ROI iv: red peaks). (c) A pictorial depiction of prostate-specific DBSI model is presented to demonstrate the association between specific prostatic histopathological structures and diffusion characteristics. Collagen fibers were associated with an anisotropic diffusion tensor, inflammatory cells are associated with highly restricted isotropic diffusion tensor, tumor cells are associated with restricted isotropic diffusion tensor, and lumen is associated with free isotropic diffusion tensor. (d) Heat maps of DBSI metric maps were generated to highlight spatial distribution of prostatic histopathological structures. ROIs = regions of interest.

### DBSI-derived diffusion metrics revealed structural complexity underlying mpMRI lesion

In a representative PCa patient, conventional mpMRI detected one hypointense lesion in both T2WI and ADC map in the transition zone (Fig. 2a). Based on the histogram, the ADC of the lesion appeared between 0.3 - 0.8 µm^2^/ms (Fig. 2b). Histologic analysis of the corresponding whole-mount prostatectomy specimen sections revealed the pathological heterogeneity within the lesion that was missed by ADC histogram (Fig. 2c). In contrast to the ADC histogram, DBSI-isotropic ADC histogram exhibited four non-overlapping distributions within the mpMRI lesion (Fig. 2d). The peak ADC of the four sub-histograms located at 0.05, 0.3, 1.5 and 3.0 µm^2^/ms, corresponding to the highly restricted (lymphocytes), restricted (tumor), hindered (stroma), and free (intra-luminal space) isotropic diffusion, respectively (Fig. 2d). Heat maps of DBSI-isotropic ADC segments corresponding to the four peaks and an anisotropic diffusion fraction were generated to reveal putative underlying cellular and structural components (Fig. 2e). A 3D-rendered DBSI-metric heat map (Fig. 2f) of prostate from a 58-year-old patient (PSA 6.49 ng/ml) localized suspicious peripheral zone PCa (pink, restricted fraction), transition zone BPH (gold, anisotropic fraction), and peripheral zone inflammation overlapping PCa (blue-green, highly-restricted fraction). These *in vivo* DBSI findings were consistent with corresponding H&E staining results of the whole-mount prostatectomy specimen sections (Fig. 2f).

**Figure 2.**
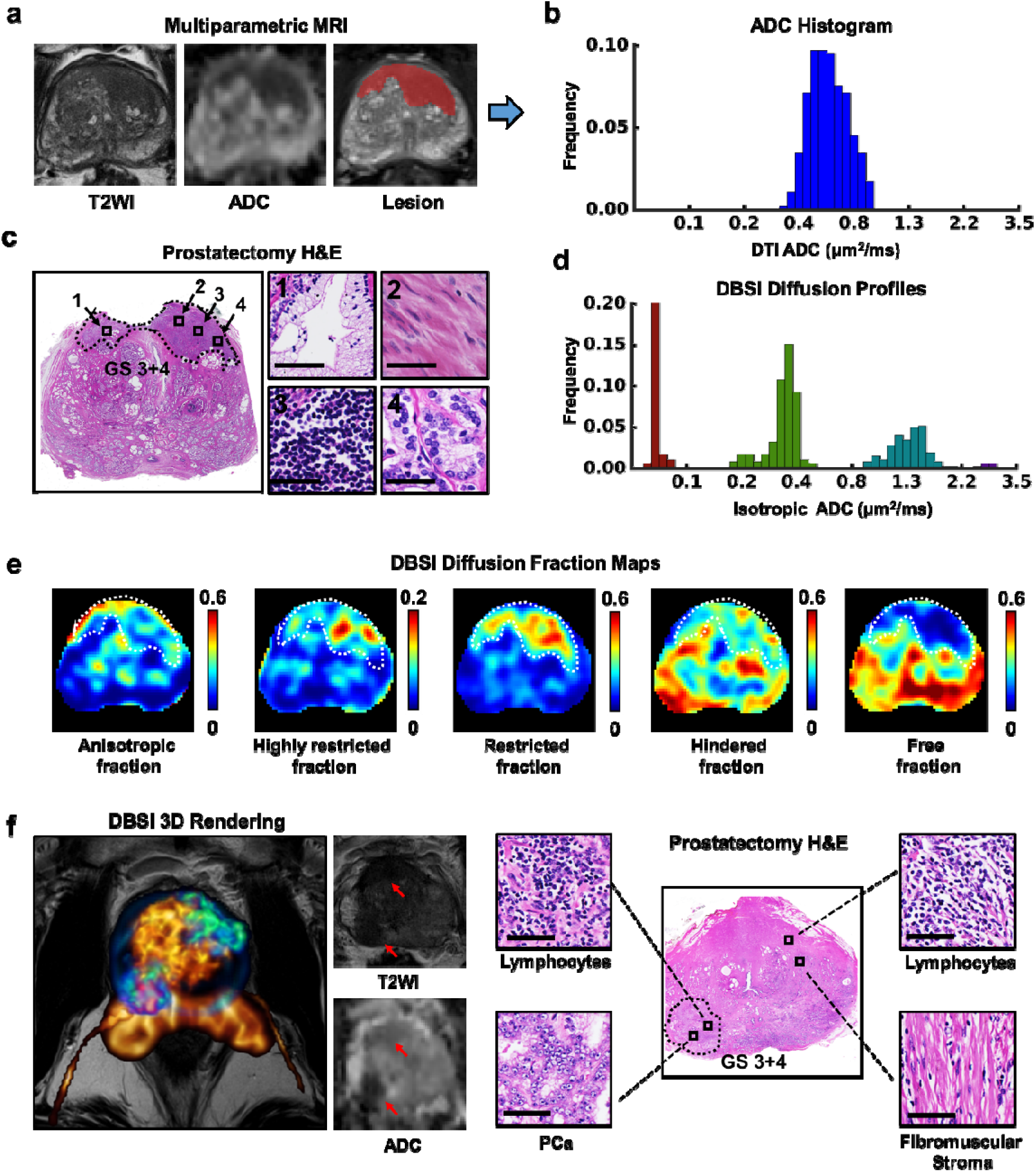
DBSI-derived diffusion metrics revealed structural complexity underlying mpMRI lesion. (a) The mpMRI images from a patient with PCa (post-op Gleason grade group 2) indicated a malignant lesion in the transition zone. (b) DTI ADC histogram analysis indicated a singular peak distribution with the range of 0.3 to 0.8 µm^2^/ms reflecting tumor presence. (c) H&E images of the whole mount section of the corresponding MRI slice also indicated a tumor region at the transition zone. The expanded view revealed the coexistence of lymphocytes, tumor cells, stromal cells, and luminal structure in the tumor region. (d) DBSI diffusion profiles exhibited four distinct groups of isotropic ADC distributions as highly restricted diffusion, restricted diffusion, hindered diffusion and free diffusion, coinciding the coexisting lymphocyte, tumor cells, stromal cells and luminal structure underlying tumor region. (e) DBSI-metric maps reflecting underlying prostatic structures and pathologies: anisotropic diffusion fraction (BPH, fibromuscular tissue), highly restricted diffusion fraction (lymphocytes), restricted diffusion fraction (cancer cells), hindered diffusion fraction (stroma), and free diffusion fraction (luminal space). (e) A 3D-rendered prostate from a patient containing PCa, inflammation, and BPH. The 3D-rendered prostate was overlaid on the T2W anatomical image. Significant BPH is visible in the transition zone (yellow channel from anisotropic fraction). Inflammation is seen in right lateral anterior peripheral zone and left lateral anterior central zones (yellow arrows; blue-green color channel imported from highly restricted fraction). PCa located at the left lateral anterior peripheral zone (purple-red channel imported from restricted fraction) surrounded by inflammatory cells. In the corresponding locations, lymphocytes, PCa and fibromuscular stroma could be identified from prostatectomy histology. T2W and ADC map showed ambiguous signals and could not provide as detailed information as DBSI did.

### Tumor cellularity correlated with DBSI-restricted fraction in *ex vivo* prostate specimen

We have demonstrated that both *ex vivo* and *in vivo* DBSI restricted isotropic diffusion fractions (restricted fraction) highly overlapped with pathologist-identified tumor lesions in prostate. To quantitatively demonstrate the relationship between DBSI restricted fraction and PCa tumor cellularity, we examined *ex vivo* DBSI metrics in histologically-identified tumor regions in five prostatectomy specimens from five different PCa patients. For comparison, conventional ADC derived from standard of care mpMRI from the same samples were also included for analysis. We performed a landmark-based thin-plate-spline (TPS) co-registration on prostatectomy specimens between MR images and H&E images to allow voxel-wise correlation of histology (H&E tumor cell counts) with ADC and DBSI-restricted fraction maps (Fig. 3a). We randomly selected ten 250 um x 250 um regions of interest (ROI) from H&E images (Fig. 3a, blue squares) and mapped them to the co-registered MRI-metric maps. Tumor cells were manually counted in each selected ROI and were correlated with the restricted fraction and ADC values in the corresponding maps. For all five specimens, restricted fraction significantly correlated with H&E quantified tumor cellularity (r^2^ = 0.78, 0.83, 0.75, 0.67 and 0.86, respectively; *p* <0.001, <0.001, 0.001, 0.004, and <0.001, respectively), while ADC exhibited inconsistent correlations with H&E findings (r^2^ = 0.01, 0.02, 0.37, 0.004 and 0.58, respectively; *p* = 0.8, 0.7, 0.06, 0.9, and 0.01, respectively) (Fig. 3b).

**Figure 3.**
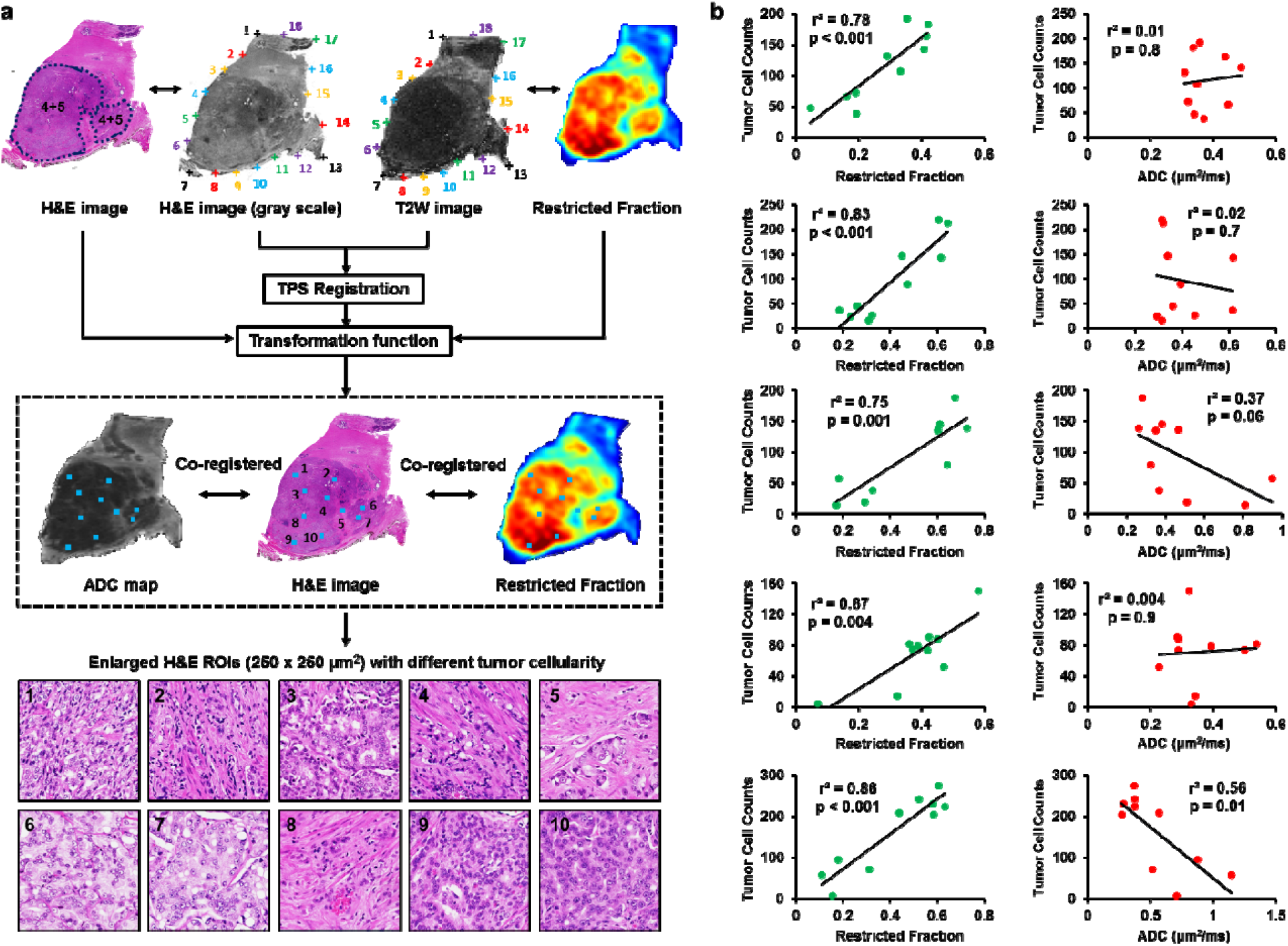
Tumor cellularity correlated with DBSI-restricted fraction but not with ADC in *ex vivo* prostate specimens. (a) We performed co-registration of T2-weighted (T2W) images and H&E images to allow voxel-wise correlation of histology-measured cellularity with ADC and DBSI-restricted fraction. Eighteen pairs of landmarks were manually placed along the perimeter of T2W images and H&E images for co-registration. The transformation function of thin-plate-spline registration was calculated and applied to warp MR images to match H&E images. Ten regions of interest (ROIs) of MRI voxel size (250 × 250 μm^2^) were randomly selected from each H&E image. The tumor cellularity in each ROI was manually counted. The ROIs were applied to corresponding co-registered MRI-metric maps for correlation and quantitative analysis. (b)Pearson’s correlation analysis indicated DBSI-restricted fraction had significant positive correlation with H&E tumor cellularity in all five specimens. However, ADC showed inconsistent correlation with tumor cellularity.

### Conventional ADC failed to distinguish PCa of different Gleason grade groups

Histogram analysis of *in vivo* ADC and DBSI-isotropic ADC from image voxels collected from 150 subjects with benign prostate (n = 54), and PCa (ranging from GG 1 – 5; n = 96) was performed and is summarized in Fig. 4. Benign peripheral and transition zone tissues were clearly distinguished from PCa by DBSI-isotropic ADC but not conventional ADC. In addition, conventional ADC failed to distinguish PCa of various Gleason scores (Figs. 4c – g, overlapping histogram). DBSI-isotropic ADC exhibited distinct profiles corresponding to benign pathologies (Figs. 4a and b) and PCa of all grades (Fig. 4c – g). Benign peripheral zone (Fig. 4a) was commonly associated with high free diffusion fraction, reflecting extensively branching luminal structures within the tissue. Benign transition zone tissues (Fig. 4b) typically exhibited various extents of hindered and free diffusion fractions. The less prominent but noticeable restricted fraction within benign transition zone tissues could potentially result from the glandular epithelial cell proliferation in epithelial BPH.

**Figure 4.**
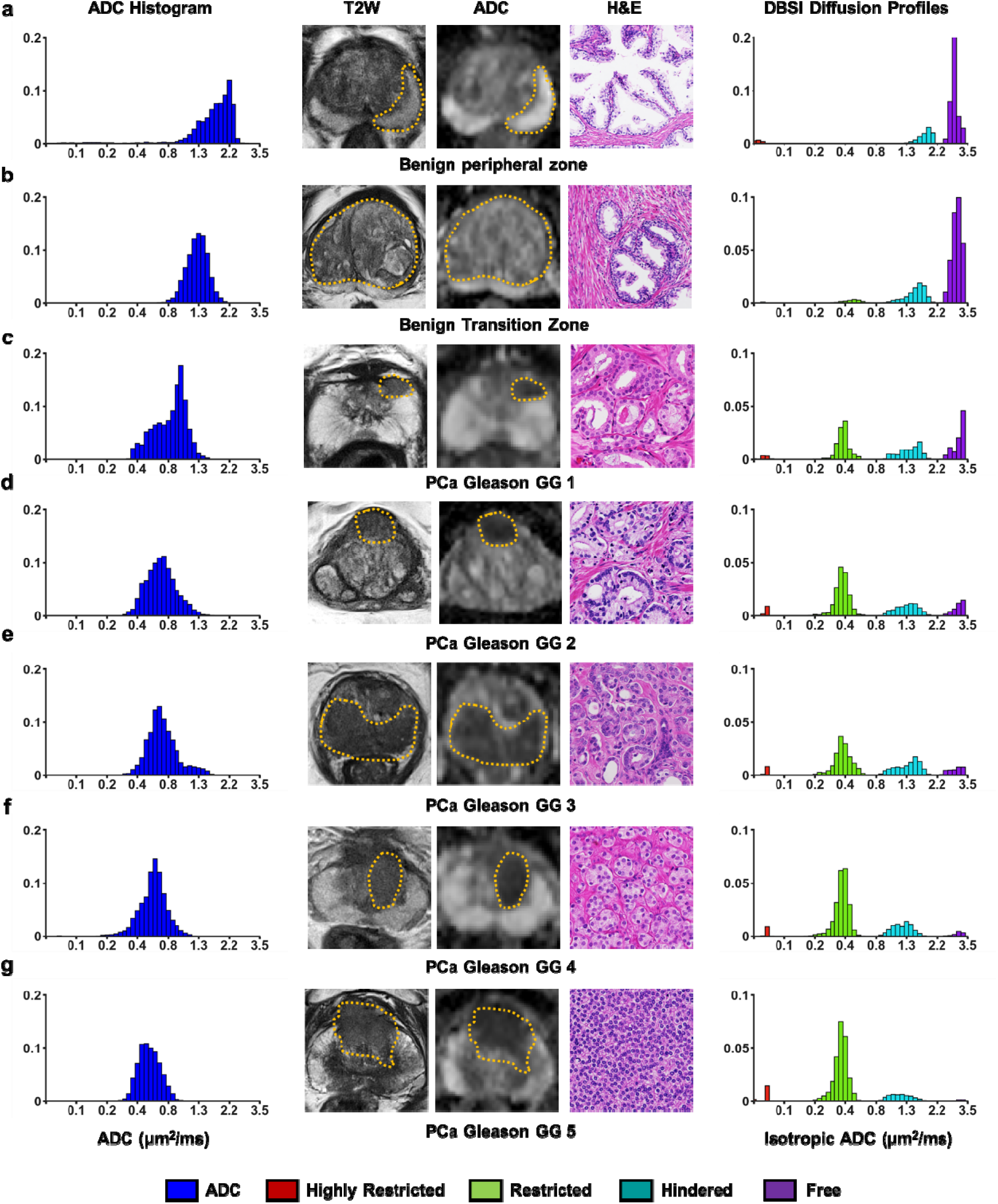
Conventional ADC failed to distinguish PCa of different Gleason grade groups. Based on histogram features, conventional ADC differentiated benign prostate (a, b) and lower-grade PCa (c, GG 1) from higher-grade PCa (d – g, GG 2 – GG 5). However, it failed to further distinguish among the higher-grade tumors. DBSI isotropic diffusivity histograms exhibited unique features for benign and PCa of all grades. Benign peripheral zone tissue (a) is associated with a high free diffusion fraction (purple) and low hindered fraction (cyan) while the benign transition zone (b) exhibited a combination of high free diffusion fraction and high hindered diffusion fraction. GG 1 tumors (c) are histologically heterogeneous, exhibiting various extents in DBSI restricted-diffusion (green), hindered-diffusion, and free-diffusion fractions. Diffusion profile of GG 3 PCa (e) had slightly increased restricted-diffusion fraction than that in GG 2 PCa (d). GG 2 and GG 3 PCa exhibited similar hindered-diffusion fraction; both grade groups have little to no signal from free-diffusion fraction, indicating an increased number of tumor/epithelial cells and the loss of luminal space. High restricted-diffusion fraction and the absence of free-diffusion fractions seen in GG 4 (f) and GG 5 tumors (g) are consistent with their histological characteristics, reflecting the hyper-cellularity in PCa with sheet-like growth pattern; few, if not the complete absence of, duct-acinar structures or stroma persists. GG = grade group.

Increased complexity of the diffusion histogram is a unique characteristic of PCa (Figs. 4c – g). In GG 1 PCa lesions (Fig. 4c), DBSI-isotropic ADC profiles exhibited highly-restricted, restricted, hindered and free isotropic diffusion fractions, reflecting the heterogeneous composition of tumor/epithelial cells, stromal cells, and luminal structures, the complexity of which is not observed in benign tissues. In contrast, the DBSI-isotropic ADC profiles exhibited decreased complexity with higher grade PCa. Profiles of GG 2 (Fig. 4d) and 3 (Fig. 4e) PCa consisted of mostly restricted and hindered isotropic diffusion signals. GG 3 tumor also exhibited increase of restricted fraction signal and a significantly decrease in free diffusion, consistent with increased tumor cellularity and loss of glandular/luminal morphology. Both GG 4 (Fig. 4f) and 5 (Fig. 4g) PCa exhibited a progressive increase in the restricted fraction component with a concomitant loss of the hindered and free isotropic diffusion fraction compared to GG 1 – 3 tumors. A noticeable trend of increasing highly restricted diffusion fraction (red, Figs. 4c – g) in GG 1 – 5 tumors, potentially reflect the increasing extents of inflammation. DBSI-isotropic ADC profiles reflected histological characteristics of GG 4 and 5 tumors, sheets of tumor cells with little to no luminal structures or stroma. The diffusion profiles modeled the structural changes underlying these tissues at different stages of malignancy uniquely, which ultimately allowed for the development of diffusion histology imaging (DHI, a pattern recognition approach using deep neural network algorithm on DBSI metrics).

### Classification of prostatic histology and Gleason scores on the voxel level

Voxel-based multi-class classification of prostatic histology using DHI resulted in a high true positive rate of PCa (0.971) and benign transition zone tissues (0.814), but low true positive rate on benign peripheral zone tissues (0.632). Significant overlap was seen between benign peripheral and transition zone tissues (Fig. 5a). We performed pairwise comparisons to examine whether PCa is distinguished from benign prostatic histology (Fig. 5a, receiver operating characteristics &precision-recall curves). DHI distinguished PCa from benign peripheral, transition zone tissues, and combination of all benign tissues (including both peripheral and transition zones) with areas under curve (AUCs) of 0.995 (95% CI: 0.995-0.996, peripheral zone), 0.985 (95% CI: 0.984-0.985, transition zone), and 0.998 (95% CI: 0.997-0.998, peripheral + transition zones), respectively (Fig. 5b). Sensitivity (> 93%) and specificity (> 94%) at optimal cut-off points (Youden’s Index) were calculated and summarized (Fig. 5b). The precision-recall (PR) analyses showed good diagnostic performances with PR-AUC > 0.98 and F_1_-scores > 0.920 (Fig. 5a).

**Figure 5.**
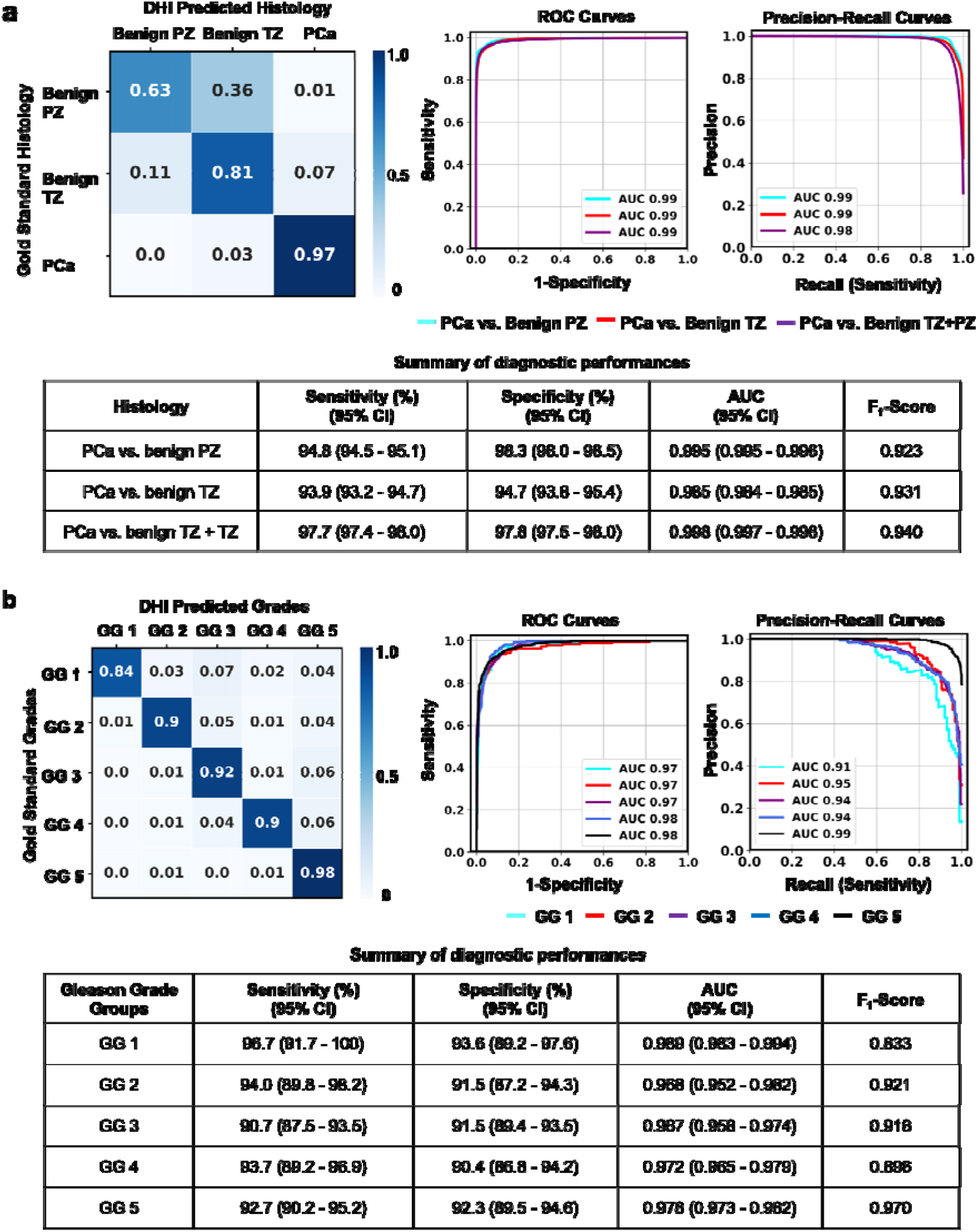
Voxel-based classification of prostatic histology and Gleason scores. (a) Confusion matrix shows concordance between DHI predictions and biopsy confirmed prostatic tissue types. PCa voxels were correctly predicted with high accuracy, but benign peripheral zone and benign transition zone voxels overlaps. ROC and precision-recall curves showed classifications with high AUC values on groups of PCa vs. benign peripheral zone (PZ), PCa vs. benign transition zone (TZ) and PCa vs. benign PZ + TZ. Table summarized AUC, sensitivity, specificity and F_1_-score values from ROC and precision-recall analyses. (b)Confusion matrix for DHI predicted PCa grade groups highly agreed with those determined histologically. ROC and precision-recall curves using one-versus-rest strategy were calculated for each Gleason grade group with high AUC values for both groups of the curves. The AUC, sensitivity, specific and F_1_-scores values of these analyses were also summarized in the table. PZ = peripheral zone. TZ = transition zone. GG = grade group. CI = confidence interval.

In the classification of Gleason scores, DHI demonstrated a 0.90 overall accuracy with recalls of 0.84, 0.90, 0.92, 0.90 and 0.98 from GG 1 through GG 5 PCa, respectively (Fig. 5b). Additionally, we used one-versus-rest strategy to assess the performances for classifying PCa grade groups. The receiver operating characteristics (ROC)-AUCs for ISUP GG 1 through GG 5 were 0.989 (95% CI: 0.983 - 0.994), 0.968 (95% CI: 0.952 - 0.982), 0.967 (95% CI: 0.958 - 0.974) and 0.978 (95% CI: 0.973 - 0.982), respectively (Fig 5b). The mean sensitivity and specificity for all of the five grade groups were above 90%. The precision-recall analysis indicated that AUC values > 0.91 for each of the five grade groups (Fig. 5b, precision-recall curves). The F_1_-scores were calculated to address class imbalance, which resulted in the respective scores of 0.833, 0.921, 0.916, 0.896 and 0.970 for GG 1 through GG 5.

### DHI noninvasively predicted PCa Gleason grade groups in patient subjects

Five representative cases with PIRADS scores of 4 and 5 were selected to test DHI model for predicting PCa and ISUP grade groups. Specifically, all image voxels from representative image slices of prostates were sent to the deep neural network DHI model for PCa/benign prediction and voxels with positive tumor prediction (pink mask) were further segmented for ISUP grade group prediction (blue mask: GG 1; light blue mask: GG 2; green mask: GG 3; yellow mask: GG 4; red mask: GG 5).

A patient underwent standard-of-care mpMRI of prostate (Fig. 6a), followed by target and systematic biopsy, and prostatectomy (Fig. S2). The conventional ADC map revealed an anterior transition zone lesion that matched the DBSI-derived restricted fraction map (Fig. 6a). DHI predicted a tumor area overlapping the anterior hyperintense DBSI restricted fraction area. For the lesion between transition and peripheral zones, DHI predicted a descrete distribution of tumor foci of ISUP GG 4 (Fig. 6a, yellow mask) and 5 (Fig. 6a, red mask). MRI/ultrasound fusion guided biopsy cores #7 and #8 PCa were GG 5 (Fig. S2), consistent with the DHI classification (Fig. 6a). The patient was considered very high risk based on the National Comprehensive Cancer Network (NCCN) risk assessment, undergoing an expedited prostatectomy after biopsy. The H&E of whole mount section of the prostatectomy specimen revealed comparable ISUP GG as seen in DHI (Fig. S2).

**Figure 6.**
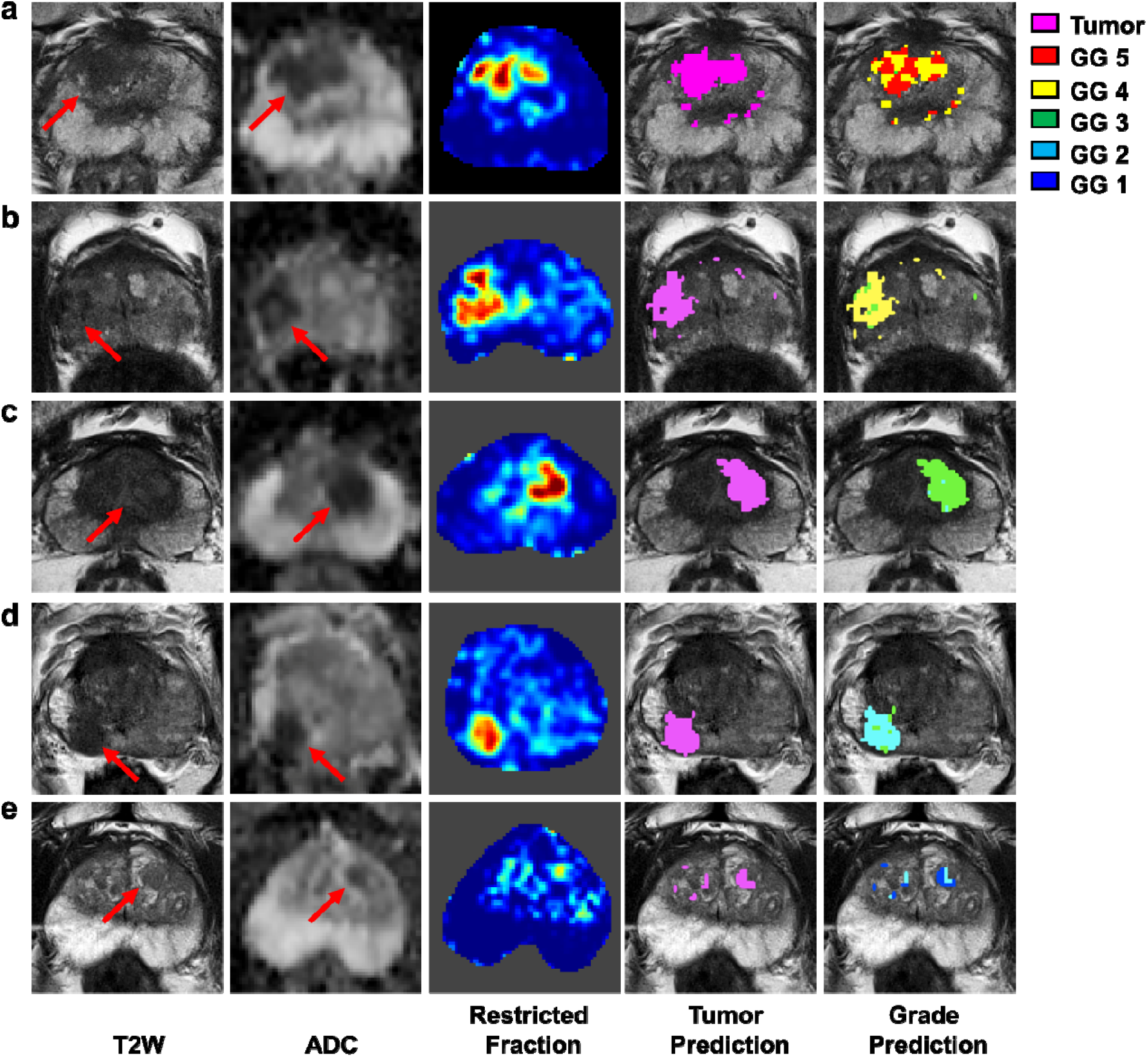
Patient-based DHI prediction of tumor and ISUP grade groups (GG). Five patients were evaluated with mpMRI and one suspicious tumor lesion was identified in each patient (red arrows in ADC maps). Patient 2 (b), 5 (e) are PI-RADS score of 4; patient 1 (a), 3 (c) and 4 (d) are PI-RADS score of 5. Arrows pointed to suspicious lesions. DHI predicted the same cancer lesions (purple masks in DHI tumor map) identified by the PI-RADS criteria. The target biopsy for all selected lesions was positive for PCa. The biopsied histology for patients 1 to 5 was GG 5, GG 4, GG 1, GG 2, and GG 1, respectively. Smaller, secondary tumor foci were also predicted for patients 1, 2 and 5. In addition to target biopsy, standard biopsy suggested positive PCa findings in multiple biopsy needle cores, indicating that some of the missed PCa lesion from mpMRI would have been detected by DHI. DHI correctly predicted the grade groups for the dominant lesions for all five patients (DHI grade maps).

In case 2, DHI predicted the majority of the lesion to be in GG 4 (Fig. 6b, yellow) with remaining voxels as GG 3 (Fig. 6b, green) voxels. The predictions of case 2 agreed with prostatectomy grade groups. Biopsy for this patient indicated 8 needle cores were PCa positive, among which 5 were GG 1, one was GG 3, and two were GG 4. For case 3, DHI predicted the primary lesion to be GG 3 (Fig. 6c, green mask) lesion with a few voxels as GG 2 (Fig. 6c, light blue mask). DHI prediction was consistent with the prostatectomy evaluation that indicated this patient to be GG 3, contradicting the targeted biopsy result that placed the lesion in GG 1 (Fig. 6c, green). In case 4, DHI predicted the primary lesion to be PCa of GG 2 (Fig. 6d, light blue mask) with a few voxels indicating GG 3 presence (Fig. 6d, green mask). This finding, again, was consistent with both targeted biopsy and prostatectomy results (GG 2). In case 5, DHI predicted the primary lesion was GG 1 (primary; Fig. 6e, blue) and GG 2 (secondary; Fig. 6e, light blue mask). Both prostatectomy and targeted biopsy indicated the lesion to be PCa of GG 1 in case 1. Two additional cases (Fig. S3a &b) were PI-RADS v2 score 4 with suspicious lesions identified in ADC maps (Fig. 3a &b, arrows). However, DHI predicted these two patients to be PCa free, in agreement with biopsy results.

## Discussion

We have demonstrated that DHI noninvasively and accurately distinguishes PCa from benign tissues reducing the false positive rate of mpMRI confounded by BPH and prostatitis signals. We also observed the specific association of DBSI derived “diffusion fingerprints” with various histopathological features of prostate glands (Fig. 2). However, the mere association between DBSI derived “diffusion fingerprints” is insufficient to unequivocally classify PCa without false positivity or accurately assess Gleason scores (Fig. 4). Thus, we employed DHI, with its deep neural network analysis of DBSI “diffusion fingerprints” as the input parameters, to accurately classify and localize PCa vs. benign tissues, and assess PCa Gleason grade groups (Fig. 4).

Normal prostate histology is rich in glandular units that are embedded in dense fibromuscular stroma (*32*). This branching duct-acinar glandular architecture allows water to diffuse more freely than in surrounding stroma or in highly cellular microenvironments (e.g. inflammation, cancer) (*32*). The transition zone consists of stromal (hindered water diffusion) and fibromuscular (anisotropic water diffusion) tissues. The periurethral muscles, anterior fibromuscular regions, and stromal BPH or fibrous tissues surrounding the BPH can all contribute to anisotropic water diffusion (*32*).

The development of PCa interrupts these glandular units and stroma structures, with increasing cellularity. Corresponding diffusion MRI features of restricted diffusion reflecting inflammatory and tumor cells are distinct from diffusion properties of benign prostate tissues. Thus, the observed association of decreased ADC with PCa is readily attributable to the increased extent of low-diffusivity epithelial cells disrupting the stroma and luminal spaces (*33*). The improved DBSI modeling of diffusion-weighted MRI signals generating unique “diffusion fingerprints” to represent underlying structural complexity and/or changes resulting from cancer presence has more successfully relfected both benign and malignant prostate tissues (Fig. 4). Thus, the combination of DBSI-”diffusion fingerprints” (i.e., structural metrics) and deep neural network algorithm has provided a noninvasive approach to differentiate Gleason grade groups of PCa of all grades (*34*).

The microstructural changes induced by PCa is complex. Glandular structures vary in PCa since some cancer cells grow between ducts sparing sufficient luminal space in contrast to other poorly differentiated cancers with more expansile masses of small and tightly packed cell clusters leaving little luminal spaces. Increased cellularity leads to the compactness of tissue and more restricted water diffusion. This lays the foundation for decreased ADC to differentiate between benign and low grade PCa from intermediate/high grade lesions (*35, 36*). ADC falls short in distinguishing intermediate from high grade PCa because ADC values of intermediate and high grade PCa significantly overlap (Fig. 4) (*35, 37*). The lack of specificity of ADC to distinguish intermediate from high grade PCa is due to the average of coexisting pathologies with opposing diffusion effects within an imaging voxel (*32*).

To improve the specificity of ADC to cancer detection, a more sophisticated diffusion MRI modeling is needed. Restriction Spectrum Imaging (RSI) is one of such attempts (*38-42*). This novel technique separates hindered and restricted diffusion to resolve a spectrum of length scales and incorporating geometric information, estimated from high-angular diffusion-weighted data based on the fiber orientation density function and 4th order spherical harmonics. Similar to DBSI, RSI also assumes a Gaussian diffusion model. RSI defined spectrum was a combination of anisotropic, restricted and free/hindered diffusion compartments (*41, 43*). RSI does not quantify ADC because it uses fiber orientation density function without directly modeling diffusion tensors as seen in DBSI. RSI assumes known free water diffusivity and a priori axial diffusivity by fixing axial diffusivity while allowing radial diffusivity to vary for modeling diffusion-weighted data, i.e., the geometrical assessment (ratio of radial/axial diffusivity) of tissue (*41, 44*). RSI has recently been applied to successfully distinguish PCa from benign tissues using cellularity index but it did not grade PCa as demonstrated herein by DHI (*42, 44*). DHI does not search through the unspecified structural or pathological features potentially embedded in the raw diffusion-weighted MRI signals as other radiomic approaches. It focuses on classification of the DBSI-derived diffusion signatures that have been well-defined with increased sensitivity and specificity to prostate structures and pathologies. DHI thus requires smaller amount of data for classifications using artificial intelligence algorithms.

DHI accurately localized and graded PCa in this proof-of-concept study suggesting that it could potentially guide biopsy and focal therapy, as well as monitoring treatment responses. The observed improvement in sensitivity and specificity of DHI to PCa diagnosis herein if validated could prevent patients with low-grade disease from undergoing unnecessary surgery, provide less invasive treatment options, stratify therapies, and monitor effectiveness of focal therapies, such as external beam radiation therapy (*45-47*) and high intensity focused ultrasound (HIFU) therapy (*48*) to spare more normal tissues surrounding the lesions. DHI could potentially be incorporated with clinical information (e.g., PSA levels, and liquid biopsy biomarkers) to improve PCa risk stratification, therefore, providing critical information on PCa prognosis and improving effectiveness of active surveillance.

In conclusion, we introduced the novel diffusion histology imaging approach, demonstrating its effectiveness as a noninvasive diagnostic tool for PCa in this small proof-of-concept study. By obtaining the DBSI-derived structural fingerprints to accurately capture the various prostate histopathological features, DHI addressed the shortcomings of standard of care mpMRI to improve detection and grading of PCa demonstrating its role as a noninvasive alternative to complement and improve the current clinical diagnosis and management of PCa.

## Materials and Methods

### Study design

Between March 2015 and August 2017, 254 patients with clinical suspicion of prostate cancer (PSA > 4.0 ng/ml) were recruited for this study. Patients who received preoperative treatments such as androgen deprivation (n = 1) or radiation therapy (n = 2) were excluded from this study. Additionally, patients with motion artifacts in image data (n = 2) and problematic image data (n = 6) were also excluded. Total 93 patients who scored lower than 3 on the Prostate Imaging– Reporting and Data System, version 2 (PI-RADS v2) criteria and with a PSA level lower than 10.0 ng/ml were considered cancer free and not biopsied. The remaining 150 patients with the PI-RADS v2 scores > 3 or PSA levels > 10 ng/ml were scheduled for combined systematic and MRI-targeted ultrasound-guided biopsy. The biopsies confirmed PCa positive in 96 patients and benign in 54 patients. Prostatectomy specimens were procured for the *ex vivo* imaging study. Prostatectomy specimens underwent high resolution MRI scanning and H&E staining. The *in vivo* patient study was approved by the local Institutional Review Board. Informed consents were waived due to the nature of retrospective study. The *ex vivo* specimen study was also approved by the local Institutional Review Board. Each patient gave informed consent before the study.

### DBSI and mpMRI data acquisition

A total of 243 patients were imaged with DBSI and mpMRI at Changhai Hospital with a 3.0-T Siemens Skyra scanner (Erlanger, Germany) with an 18-channel pelvic phased-array coil. The DBSI protocol involved acquiring diffusion-weighted echo-planar imaging data with the following data acquisition parameters: repetition time (TR) 5000 ms, echo time (TE) 88 ms, 4 averages, FOV 112 x 140 mm^2^, in-plane resolution 2 x 2 mm^2^, 24 slices at 4-mm thick, 25-dir icosahedral diffusion encoding scheme with maximum b-value 1500 s/mm^2^. Standard mpMRI protocol, including T1-weighted, T2-weighted, diffusion-weighted imaging and dynamic contrast-enhanced T1-weighted sequences, was performed on all patients. Prostate Imaging Radiology Assessment and Diagnostic System Version 2 (PI-RADS v2) was used by experienced radiologists for assessing suspicious prostate cancer on mpMRI.

### Ultrasound-guided target and systematic biopsy

Multi-parametric MRI suspicious of PCa was scored by an experienced radiologist according to PI-RADS v2. A total of 150 patients with PI-RADS v2 score higher than 3 underwent transperineal ultrasound (TPUS)-guided systematic biopsy, and TPUS-MRI-fusion guided biopsy. Fusion of MRI with the real-time transperineal ultrasound image of the prostate during biopsy was performed by either cognitive or software-facilitated registration. Biopsy-obtained tissues were analyzed and graded by clinical pathologists. International Society of Urological Pathology (ISUP) grading of prostate cancer (*49*) was used to assess the cancer lesions.

### Prostatectomy specimen preparation for *ex vivo* MRI

After prostatectomy, each prostate specimen was immediately fixed with 10% formalin in phosphate buffered saline (PBS, PH = 7.4) for at least 10 hours. The specimen was step-sectioned at 4-mm intervals from base to apex.

### *Ex vivo* MRI of prostatectomy specimen

Prostatectomy specimens were formalin-fixed at time of collection and examined using a 4.7-T Agilent DirectDrive^*™*^ small-animal MRI system (Agilent Technologies, Santa Clara, CA) equipped with a Magnex/Agilent HD imaging gradient coil (Magnex/Agilent, Oxford, UK) capable of pulsed-gradient strengths of up to 58 G/cm and a gradient rise time ≤ 295 µs. A multi-echo spin-echo diffusion weighting sequence employing 25 diffusion-encoding directions with maximum b-value = 3000 s/mm^2^ was employed to acquire diffusion-weighted images. The imaging parameters were: TR 1500 ms, TE 40 ms, time between application of gradient pulses 20 ms, diffusion gradient on time 8 ms, slice thickness 0.5 mm, field-of-view 24 × 24 mm^2^, data matrix 96 × 96, number of average 1. Total acquisition time was approximately 1 hour and 20 minutes. T1W, T2W and DW images with the same image resolution were also acquired.

### Histological analysis

Prostatectomy specimens were embedded in paraffin and underwent sequential sectioning at 5-μm thick after MRI. Sections were individually stained with hematoxylin and eosin (H&E). Histology slides were digitized using NanoZoomer 2.0-HT System (Hamamatsu, Japan) with a 20× objective for analyses.

### Co-registration between *ex vivo* MRI and histology images

Two-dimensional (2D) thin plate spline (TPS) registration was performed using *MIPAV* Version 10.0.0 (NIH, Bethda, MD) as described previously (*28*) to co-register *ex vivo* MR images with histology images. We first ensure the plane of histology section of the prostate specimens matched closely with the slice plane of the corresponding T2W images. Then the RGB H&E images were converted to grayscale using the Pillow package in Python (https://pillow.readthedocs.io/en/3.1.x/index.html). After these preprocessing steps, around eighteen pairs of landmarks along the perimeter of each specimen were manually placed on the gray scale H&E images as well as the T2W images (inherently co-registered with ADC and DBSI maps) to compute the transformation matrix for matching H&E images with MRI. After successful image co-registration, ten regions with the same size of MRI voxels (250 × 250 μm^2^) were randomly selected from the H&E images to correlate the tumor cell counts in each region with corresponding ADC and restricted fraction values (Fig. 3a).

### Diffusion basis spectrum imaging

We developed diffusion basis spectrum imaging (DBSI) (*24*) to model DWI signals of PCa as a linear combination of an anisotropic diffusion tensor and a spectrum of isotropic diffusion tensors:

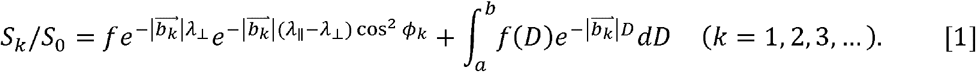

The quantities *S*_0_, *S*_*k*_ and 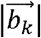 are the proton-density-weighted signal, diffusion-weighted signal, and *b*-value of the *k*^*th*^ diffusion gradient, ϕ_*k*_ is the angle between the *k*^*th*^ diffusion gradient and the principal direction of the fiber-like structures, λ_||_ and λ_┴_ are axial (AD) and radial (RD) diffusivity of the fiber-like structures (may reflect the compactness of the structure), *f* is the signal intensity fraction for fibers (representing fiber density, e.g., benign prostatic hyperplasia (BPH), in the image voxel), and *a* and *b* are the low and high diffusivity limits for the isotropic diffusion spectrum, *f*(*D*), containing contributions from cellularity and luminal water.

DBSI was initially developed to assess CNS pathologies. It has been successfully applied in animal models and translated to various CNS diseases (*24, 27, 50*). Although these CNS related interpretations do not apply to prostate, we have performed a supervised DBSI signal processing to readjust *diffusion tensor basis sets* (a dictionary of varied anisotropic and isotropic diffusion tensors distributed in q-space for modeling DWI signal) referencing histological findings from radical prostatectomy and biopsy specimens. Based on these preliminary results, we have observed that DBSI-derived anisotropic fraction is highly associated with BPH and some PCa regions, reflecting fibromuscular fiber density in each image voxel. The other hallmark prostate histopathology of lymphocytes, PCa, stromal tissues and luminal structures exhibits high correlation with highly-restricted diffusion (0 – 0.1 μm^2^/ms ∼ *lymphocytes*), restricted diffusion (0.1 – 0.8 μm^2^/ms ∼ *PCa*), hindered diffusion (0.8 – 2.5 μm^2^/ms ∼ *stromal tissues*) and free diffusion (2.5 – 3.0 μm^2^/ms ∼ *luminal water*), respectively.

The main difference between *in vivo* and *ex vivo* diffusion MRI is the tissue temperature. In general, *in vivo*/*ex vivo* apparent diffusion coefficients differ by a 3:2 ratio based on temperature difference. The temperature effect is masked by the diffusion restriction resulting from the size of cells and diffusion times. In our Monte-Carlo simulation of restricted diffusion in cells of diameter ranging between 5 - 20 µm, we observed no differences in ADC of cells between *in vivo* (at 37°C) and *ex vivo* (at 20°C). Thus, *in vivo* ADC thresholds for lymphocytes, tumor cells, stroma, and lumen can be established using high-quality *ex vivo* data matched by the histology defined tissues types, i.e., the gold standard. BPH definition is straightforward for DBSI analysis since the axial and radial diffusivity of the anisotropic diffusion tensor are allowed to change during the modeling. Thus, the *in vivo* and *ex vivo* difference in anisotropic diffusion tensor is not significant, as supported by our previous CNS studies on autopsy specimens.

### Image processing

Voxel-based DTI and DBSI analyses were performed by an in-house software developed using MATLAB^®^ (MathWorks; Natick, MA). Cancer regions of interest (ROIs) were delineated by one experienced radiologist (Q.Y.) on mpMRI by referencing pathologist-marked tumor regions on histopathological images of prostatectomy specimens using ITK-SNAP (http://www.itksnap.org/). For patients without radical prostatectomy, ROIs were defined on mpMRI regions with positive target biopsies. Gleason score of each lesion was evaluated from corresponding histopathological whole mount section images (n = 92) or targeted biopsies (n = 4). For PCa negative cohorts, representative image slices were chosen to draw ROIs for benign peripheral and transition zones. The ROIs were then applied to DBSI maps for further analysis.

### DNN model development and optimization

We constructed the DNN models using TensorFlow 2.0 frameworks (*51*) in Python. DNN models with various numbers of hidden layers, nodes and training epochs were tested to optimize the model. Afterwards, the DNN model was composed of 10 fully connected hidden layers. Batch normalization was performed with a mini-batch size of 200 before feeding data into the next hidden layer to optimize the model and to prevent overfitting. We employed exponential linear units (ELU) to activate specific functions in each hidden layer. The final layer was a fully connected softmax layer that produces a likelihood distribution over the output classes. We trained the network with random initialization of the weights as described in He et al (*52*). The Adam optimizer was used with the default parameters of β_1_ = 0.9 and β _2_= 0.999 and a mini-batch size of 200. The learning rate was manually tuned to achieve the fastest convergence (1 × 10^−3^). We chose cross-entropy loss function and trained the model to minimize the error rate on the development dataset. In general, the hyper-parameters of the neural network architecture and optimization algorithm were chosen through a combination of grid search and manual tuning.

For DNN modeling of PCa vs. benign tissues, the training dataset was built with image voxels (n = 488,762) from 93 PI-RADS v2. Score < 3 patients and 60 biopsy-confirmed PCa positive patients. Image voxels (n = 275,774) from 52 patients with biopsy-confirmed PCa negative and 30 patients with biopsy-confirmed PCa positive were used to construct tuning dataset and test dataset with 1:1 ratio. We performed a multi-class classification and binary classifications (PCa vs. Benign PZ, PCa vs. benign TZ, and PCa vs. all benign tissues) among benign PZ, benign TZ and PCa using these datasets. For classification of ISUP Gleason grade groups, we used 23,327 image voxels from 90 biopsy-confirmed PCa patients, randomly grouping these voxels into training, tuning, and test datasets with 8:1:1 ratio. To balance data from different grade groups, we applied a synthetic minority oversampling technique (SMOTE) (*53*) to over-sample the minority group by introducing synthetic feature samples. This data balancing approach has been demonstrated to be beneficial for avoiding over-fitting and improving model generalization (*53, 54*). Data balancing were only applied to the training dataset, while the validation and test dataset was kept unchanged. We performed a multi-class classification on ISUP grade groups 1 through 5, followed by binary classifications using one-versus-rest strategy for each grade group.

To further validate the model performances on predicting PCa and ISUP grade group at individual patient level, we tested 5 biopsy-confirmed PCa positive and two biopsy-confirmed PCa negative (PIRADS scores ≥ 4) patients. The predicted results were compared with the histopathological evaluations from biopsy and prostatectomy sections. Specifically, voxels of the representative image slice of prostate were sent to the above model for binary prediction on PCa. Subsequently, image voxels with positive PCa were further predicted for ISUP grade group.

The diffusion metrics used for DNN modeling of PCa vs. benign tissues includes: isotropic ADC, anisotropic fraction, anisotropic tensor fractional anisotropy (FA), anisotropic tensor axial diffusivity (AD), anisotropic tensor radial diffusivity (RD), highly restricted isotropic fraction, highly restricted isotropic diffusivity, restricted fraction, restricted isotropic diffusivity, hindered isotropic diffusion fraction (hindered fraction), hindered isotropic diffusivity, free isotropic diffusion fraction (free fraction), and free isotropic diffusivity. For DNN modelling of classification of Gleason grade groups, we also incorporated diffusion metrics from DTI modelling, including mean ADC, FA, AD and RD.

### Statistical analysis

We used Pearson correlation to assess the relationship between histology and MRI measurements. Statistically significant results were determined at a pre-determined alpha level of 0.05. We derived confusion matrices to demonstrate the agreement between DHI predictions and respective prostatic histology and PCa grade groups assessed by the gold standard histology. The one-versus-rest strategy was implemented for binary classification to assess model discrimination amongst respective prostatic histology or ISUP grade groups. We performed receiver operating characteristics (ROC) analysis and calculated area under curves (AUC) to assess sensitivity and specificity at the optimal cut off point (Youden Index) (*55*). The precision-recall curves and F_1_-scores were calculated to demonstrate the relationship between precision and recall, which provides complementary information to the ROC curve, as the dataset included imbalanced classes. All values bounding the 95% confidence intervals were calculated with bootstrapping methods iterated 10,000 times (*56*). Statistical metrics and curves were calculated by packages from Scikit-learn (*57*).

## Supplementary list

Table S1. Patient information.

Fig. S1. Flowchart of patient recruitment process.

Fig. S2. Representative case with transperineal biopsy and prostatectomy whole mount section. Fig. S3. False positive cases from mpMRI were correctly predicted by DHI.

## Author contributions

S.K.S., J.L., Z.Y., Q.Y. and J.I., initiated the project and collaboration, and drafted research design. Q.Y., C.S., P.S. and L.C. designed the clinical setup. Q.Y., Y.Z., L.G., Z.W., M.W., and Y.C. acquired the data. Y.Z., Q.Y. and Y.Y. created the dataset and defined clinical labels. Z.Y., J.L., P.S., A.T.W., R.Y., C.S., A.G., J.Z., S.E.G., and J.D.V. analyzed the data. Z.Y. and A.T.W. developed the network architectures, training and testing setup. Z.Y., J.L., Q.Y., J.E.I., E.H.K., Q.Y. and S.K.S. wrote the paper. All authors have reviewed and approved the final version of the manuscript.

## Data and materials availability

All the data supporting the findings of this study are available from the corresponding authors upon reasonable request.

## References and notes

1. R. L. Siegel, K. D. Miller, A. Jemal, Cancer statistics, 2020. CA Cancer J Clin 70, 7–30 (2020).

2. X. Filella, L. Foj, Prostate Cancer Detection and Prognosis: From Prostate Specific Antigen (PSA) to Exosomal Biomarkers. Int J Mol Sci 17, (2016).

3. D. M. Berney et al., Validation of a contemporary prostate cancer grading system using prostate cancer death as outcome. Br J Cancer, (2016).

4. M. Wenzel et al., Complication Rates After TRUS Guided Transrectal Systematic and MRI-Targeted Prostate Biopsies in a High-Risk Region for Antibiotic Resistances. Front Surg 7, 7 (2020).

5. P. F. Pinsky, H. L. Parnes, G. Andriole, Mortality and complications after prostate biopsy in the Prostate, Lung, Colorectal and Ovarian Cancer Screening (PLCO) trial. BJU Int 113, 254–259 (2014).

6. D. Olvera-Posada et al., A Population-Based Cohort Study of the Impact of Infectious Complications Requiring Hospitalization after Prostate Biopsy on Radical Prostatectomy Surgical Outcomes. Urology 121, 139–146 (2018).

7. F. H. Schroder et al., Screening and prostate cancer mortality: results of the European Randomised Study of Screening for Prostate Cancer (ERSPC) at 13 years of follow-up. Lancet 384, 2027–2035 (2014).

8. F. H. Schroder et al., Screening and prostate-cancer mortality in a randomized European study. N Engl J Med 360, 1320–1328 (2009).

9. B. Ehdaie et al., The impact of repeat biopsies on infectious complications in men with prostate cancer on active surveillance. J Urol 191, 660–664 (2014).

10. L. P. Bokhorst et al., Complications after prostate biopsies in men on active surveillance and its effects on receiving further biopsies in the Prostate cancer Research International: Active Surveillance (PRIAS) study. BJU Int 118, 366–371 (2016).

11. H. U. Ahmed et al., Diagnostic accuracy of multi-parametric MRI and TRUS biopsy in prostate cancer (PROMIS): a paired validating confirmatory study. Lancet 389, 815–822.

12. J. L. Leake et al., Prostate MRI: access to and current practice of prostate MRI in the United States. J Am Coll Radiol 11, 156–160 (2014).

13. A. R. Padhani et al., PI-RADS Steering Committee: The PI-RADS Multiparametric MRI and MRI-directed Biopsy Pathway. Radiology 292, 464–474 (2019).

14. J. C. Weinreb et al., PI-RADS Prostate Imaging - Reporting and Data System: 2015, Version 2. Eur Urol 69, 16–40 (2016).

15. K. Sklinda, B. Mruk, J. Walecki, Active Surveillance of Prostate Cancer Using Multiparametric Magnetic Resonance Imaging: A Review of the Current Role and Future Perspectives. Med Sci Monit 26, e920252 (2020).

16. R. Jayadevan et al., Magnetic Resonance Imaging-Guided Confirmatory Biopsy for Initiating Active Surveillance of Prostate Cancer. JAMA Netw Open 2, e1911019 (2019).

17. A. B. Rosenkrantz, S. S. Taneja, Radiologist, be aware: ten pitfalls that confound the interpretation of multiparametric prostate MRI. AJR Am J Roentgenol 202, 109–120 (2014).

18. A. B. Rosenkrantz, A. Oto, B. Turkbey, A. C. Westphalen, Prostate Imaging Reporting and Data System (PI-RADS), Version 2: A Critical Look. AJR Am J Roentgenol 206, 1179–1183 (2016).

19. N. A. Pickersgill et al., Accuracy and Variability of Prostate Multiparametric Magnetic Resonance Imaging Interpretation Using the Prostate Imaging Reporting and Data System: A Blinded Comparison of Radiologists. Eur Urol Focus 6, 267–272 (2020).

20. N. A. Pickersgill et al., The Accuracy of Prostate Magnetic Resonance Imaging Interpretation: Impact of the Individual Radiologist and Clinical Factors. Urology 127, 68–73 (2019).

21. R. S. Wang et al., Determination of the Role of Negative Magnetic Resonance Imaging of the Prostate in Clinical Practice: Is Biopsy Still Necessary? Urology 102, 190–197 (2017).

22. A. Amin et al., The Magnetic Resonance Imaging in Active Surveillance (MRIAS) Trial: Use of Baseline Multiparametric Magnetic Resonance Imaging and Saturation Biopsy to Reduce the Frequency of Surveillance Prostate Biopsies. J Urol 203, 910–917 (2020).

23. G. N. Tran et al., Magnetic Resonance Imaging-Ultrasound Fusion Biopsy During Prostate Cancer Active Surveillance. Eur Urol 72, 275–281 (2017).

24. Y. Wang et al., Quantification of increased cellularity during inflammatory demyelination. Brain : a journal of neurology 134, 3590–3601 (2011).

25. T. H. Lin et al., Noninvasive quantification of axonal loss in the presence of tissue swelling in traumatic spinal cord injury mice. J Neurotrauma, (2018).

26. T. H. Lin et al., Diffusion MRI quantifies early axonal loss in the presence of nerve swelling. Journal of neuroinflammation 14, 78 (2017).

27. Y. Wang et al., Differentiation and quantification of inflammation, demyelination and axon injury or loss in multiple sclerosis. Brain : a journal of neurology 138, 1223–1238 (2015).

28. Z. Ye et al., Diffusion Histology Imaging Combining Diffusion Basis Spectrum Imaging (DBSI) and Machine Learning Improves Detection and Classification of Glioblastoma Pathology. Clin Cancer Res, (2020).

29. P. Sun et al., Diffusion basis spectrum imaging provides insights into MS pathology. Neurology -Neuroimmunology Neuroinflammation 7, e655 (2020).

30. C. Cortes, V. Vapnik, Support-vector networks. Machine Learning 20, 273–297 (1995).

31. C.-C. Chang, C.-J. Lin, LIBSVM: A library for support vector machines. ACM Transactions on Intelligent Systems and Technology 2, 27 (2011).

32. J. Xu et al., Magnetic resonance diffusion characteristics of histologically defined prostate cancer in humans. Magnetic Resonance in Medicine 61, 842–850 (2009).

33. A. Chatterjee et al., Changes in Epithelium, Stroma, and Lumen Space Correlate More Strongly with Gleason Pattern and Are Stronger Predictors of Prostate ADC Changes than Cellularity Metrics. Radiology 277, 751–762 (2015).

34. D. F. Gleason, G. T. Mellinger, Prediction of Prognosis for Prostatic Adenocarcinoma by Combined Histological Grading and Clinical Staging. Journal of Urology 111, 58–64 (1974).

35. T. Hambrock et al., Relationship between Apparent Diffusion Coefficients at 3.0-T MR Imaging and Gleason Grade in Peripheral Zone Prostate Cancer. Radiology 259, 453–461 (2011).

36. O. F. Donati et al., Prostate Cancer Aggressiveness: Assessment with Whole-Lesion Histogram Analysis of the Apparent Diffusion Coefficient. Radiology 271, 143–152 (2014).

37. S. Woo, S. Y. Kim, J. Y. Cho, S. H. Kim, Preoperative Evaluation of Prostate Cancer Aggressiveness: Using ADC and ADC Ratio in Determining Gleason Score. American Journal of Roentgenology 207, 114–120 (2016).

38. R. L. Brunsing et al., Restriction spectrum imaging: An evolving imaging biomarker in prostate MRI. Journal of Magnetic Resonance Imaging 45, 323–336 (2017).

39. K. C. McCammack et al., In vivo prostate cancer detection and grading using restriction spectrum imaging-MRI. Prostate Cancer and Prostatic Diseases 19, 168–173 (2016).

40. A. Latysheva et al., Diagnostic utility of Restriction Spectrum Imaging in the characterization of the peritumoral brain zone in glioblastoma: Analysis of overall and progression-free survival. Eur J Radiol 132, 109289 (2020).

41. N. S. White et al., Diffusion-weighted imaging in cancer: physical foundations and applications of restriction spectrum imaging. Cancer Res 74, 4638–4652 (2014).

42. G. Yamin et al., Voxel Level Radiologic-Pathologic Validation of Restriction Spectrum Imaging Cellularity Index with Gleason Grade in Prostate Cancer. Clin Cancer Res 22, 2668–2674 (2016).

43. N. S. White, T. B. Leergaard, H. D’Arceuil, J. G. Bjaalie, A. M. Dale, Probing tissue microstructure with restriction spectrum imaging: Histological and theoretical validation. Hum Brain Mapp 34, 327–346 (2013).

44. M. A. Liss et al., MRI-Derived Restriction Spectrum Imaging Cellularity Index is Associated with High Grade Prostate Cancer on Radical Prostatectomy Specimens. Front Oncol 5, 30 (2015).

45. T. M. Pisansky, External-beam radiotherapy for localized prostate cancer. N Engl J Med 355, 1583–1591 (2006).

46. M. J. Zelefsky et al., High dose radiation delivered by intensity modulated conformal radiotherapy improves the outcome of localized prostate cancer. J Urol 166, 876–881 (2001).

47. E. S. Wisenbaugh et al., Proton beam therapy for localized prostate cancer 101: basics, controversies, and facts. Rev Urol 16, 67–75 (2014).

48. E. Baco et al., Hemi salvage high-intensity focused ultrasound (HIFU) in unilateral radiorecurrent prostate cancer: a prospective two-centre study. BJU International 114, 532–540 (2014).

49. J. I. Epstein et al., The 2014 International Society of Urological Pathology (ISUP) Consensus Conference on Gleason Grading of Prostatic Carcinoma. The American Journal of Surgical Pathology 40, 244–252 (2016).

50. C. W. Chiang et al., Quantifying white matter tract diffusion parameters in the presence of increased extra-fiber cellularity and vasogenic edema. NeuroImage 101, 310–319 (2014).

51. Mart et al., paper presented at the Proceedings of the 12th USENIX conference on Operating Systems Design and Implementation, Savannah, GA, USA, 2016.

52. K. He, X. Zhang, S. Ren, J. Sun, in 2015 IEEE International Conference on Computer Vision (ICCV). (2015), pp. 1026–1034.

53. N. V. Chawla, K. W. Bowyer, L. O. Hall, W. P. Kegelmeyer, SMOTE: Synthetic minority over-sampling technique. J Artif Intell Res 16, 321–357 (2002).

54. H. Haibo, B. Yang, E. A. Garcia, L. Shutao, in 2008 IEEE International Joint Conference on Neural Networks (IEEE World Congress on Computational Intelligence). (2008), pp. 1322–1328.

55. M. D. Ruopp, N. J. Perkins, B. W. Whitcomb, E. F. Schisterman, Youden index and optimal cut-point estimated from observations affected by a lower limit of detection. Biometrical J 50, 419–430 (2008).

56. J. Carpenter, J. Bithell, Bootstrap confidence intervals: when, which, what? A practical guide for medical statisticians. Stat Med 19, 1141–1164 (2000).

57. F. Pedregosa et al., Scikit-learn: Machine Learning in Python. J Mach Learn Res 12, 2825–2830 (2011).

