## supplement for "Deep neural network analysis employing diffusion basis spectrum imaging metrics as classifiers improves prostate cancer detection and grading"

**Supplemental figures and table**

### **Supplementary table 1.** Patient information.

| **Table 1. Patient Characteristics** | | |
| --- | --- | --- |
| **Total Patients (n = 243)** |  | |
| Age (yr), mean (SD) | 65.8 (9.7) | |
| PSA Level (ng/ml), mean (SD) | 25.4 (12.3) | |
| **Patients with PIRADS score ≤ 3 (n = 96)** |  | |
| Age (yr), mean (SD**)** | 62.0 (10.7) | |
| PSA Level (ng/ml), mean (SD) | 9.8 (3.9) | |
| **Patients with PCa negative biopsy (n = 54)** |  | |
| Age (yr), mean (SD) | 65.0 (8.2) | |
| PSA Level (ng/ml), mean (SD) | 11.5 (9.6) | |
| **Patients with PCa positive biopsy (n = 93)** |  | |
| Age (yr), mean (SD) | 69.3 (8.1) | |
| PSA Level (ng/ml), mean (SD) | 27.9 (39.0) | |
| **Biopsy Gleason Scores (n = 391), No. (% of total)** | | |
| Grade group 1 (Gleason score 3 + 3 = 6) | | 92 (23.5) |
| Grade group 2 (Gleason score 3 + 4 = 7) | | 65 (16.6) |
| Grade group 3 (Gleason score 4 + 3 = 7) | | 65 (16.6) |
| Grade group 4 (Gleason score 8) | | 72 (18.4) |
| Grade group 5 (Gleason score 9 - 10) | | 97 (24.7) |
| **Pathologic PCa Staging (n = 92), No. (% of total)** | |  |
| T2a | | 14 (15.2) |
| T2b | | 2 (2.5) |
| T2c | | 23 (25.0) |
| T3a | | 27 (29.3) |
| T3b | | 19 (23.4) |
| T4 | | 7 (7.6) |
| **Cancer Risk Stratification (NCCN Guideline), No. (% of total)** | | |
| Low | 2 (2.6) | |
| Intermedia | 17 (22.4) | |
| High | 35 (46.1) | |
| Very High | 22 (28.9) | |
| **Capsular Invasion, No. (% of total)** | 33 (62.3) | |
| **Seminal Invasion, No. (% of total)** | 10 (18.9) | |
| **Lymph Node Metastasis, No. (% of total)** | 6 (11.3) | |
| **Nerve Invasion, No. (% of total)** | 32 (60.4) | |

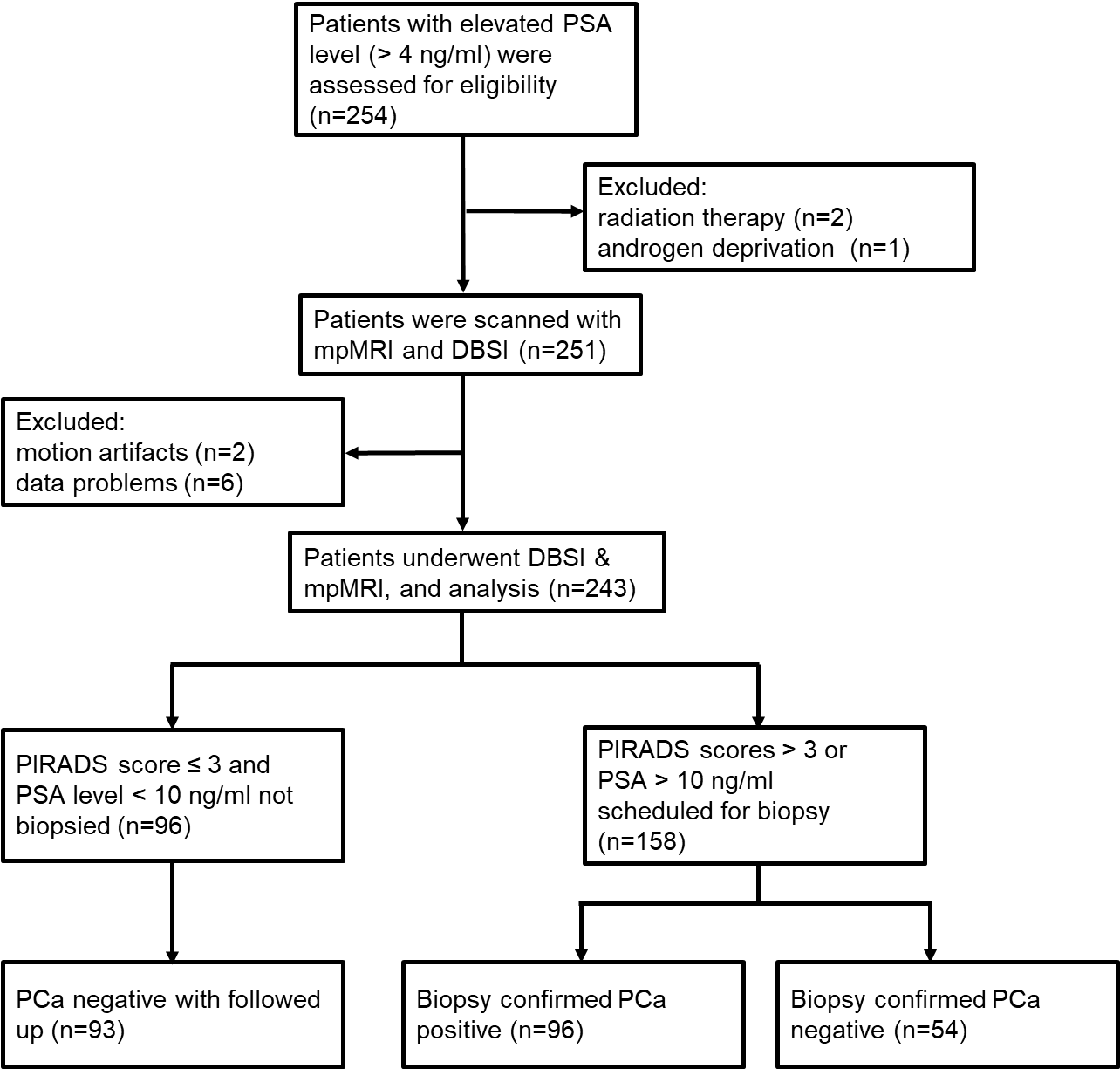

### **Figure S1.** Flowchart of patient recruitment process.

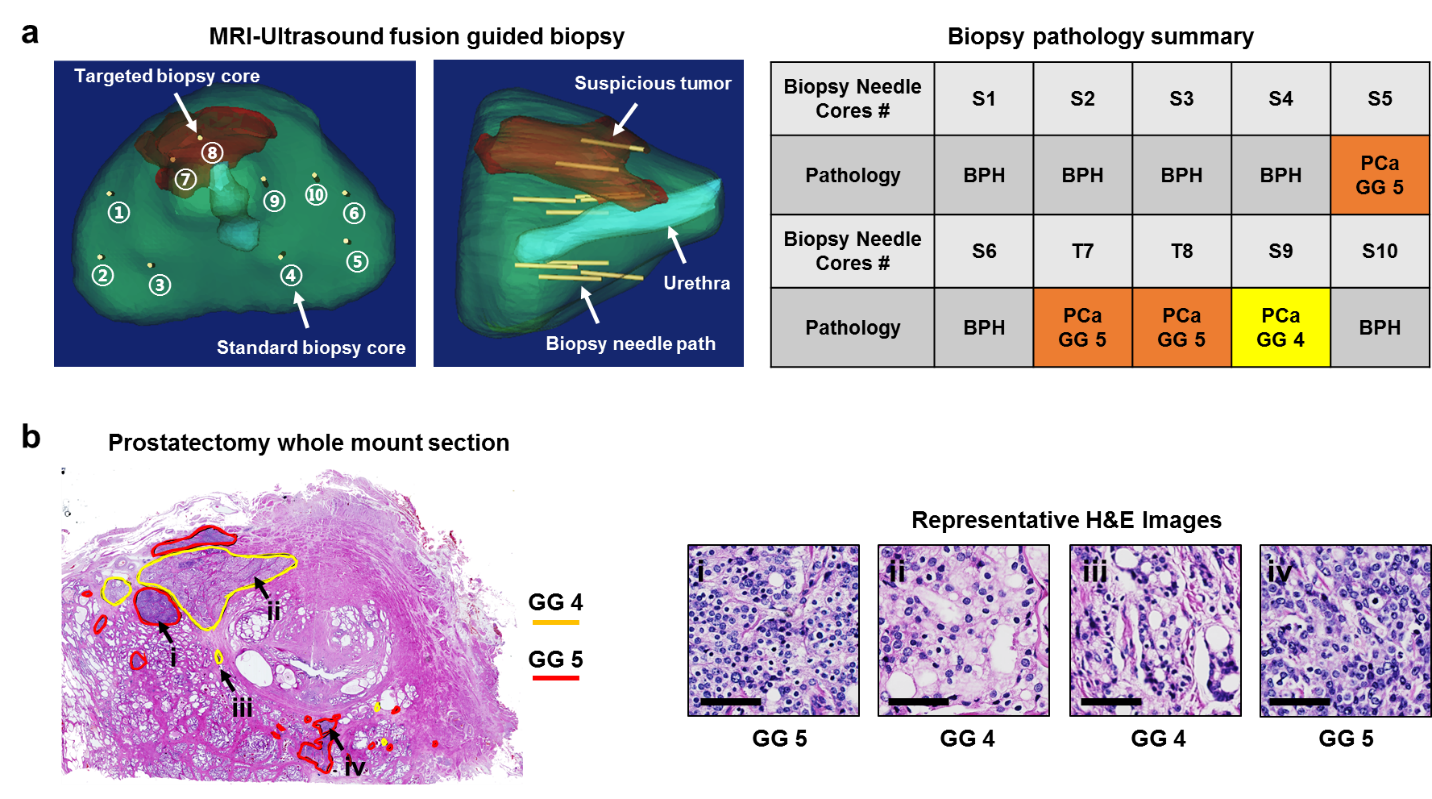

**Figure S2. Representative case with transperineal biopsy and prostatectomy whole mount section. (a)** One representative patient underwent mpMRI, DBSI and DHI analyses. MRI/ultrasound fusion guided biopsy was employed to perform transperineal targeted and systematic biopsy. Suspicious tumor region was rendered from mpMRI for targeted biosy (a, transverse view & saggital view; brown area). Two targeted biopsy (b, tranverse view; T7 & T8) and eight systematic biopsy needle cores (a, tranverse view; S1 – S6, S9 & S10) were performed and the corresponding histopathology were evaluated (a, biopsy pathology summary). Biopsy core T7, T8 and S9 agreed with the DHI’s predictions on tumor and ISUP grade groups (a, DHI). However, PCa with GG 5 from systematic biopsy core S5 was missed by DHI. After prostatectomy, whole mount section specimen was obtained and stained with H&E (b). Tumor regions were identified and graded with yellow (GG 4) and red (GG 5) outlines. Represenatative tumor regions from selected tumor foci were enlarged to show the detailed histologic characteristics of GG4 (c: i, iv) and GG5 (c: ii, iii). GG = grade group.

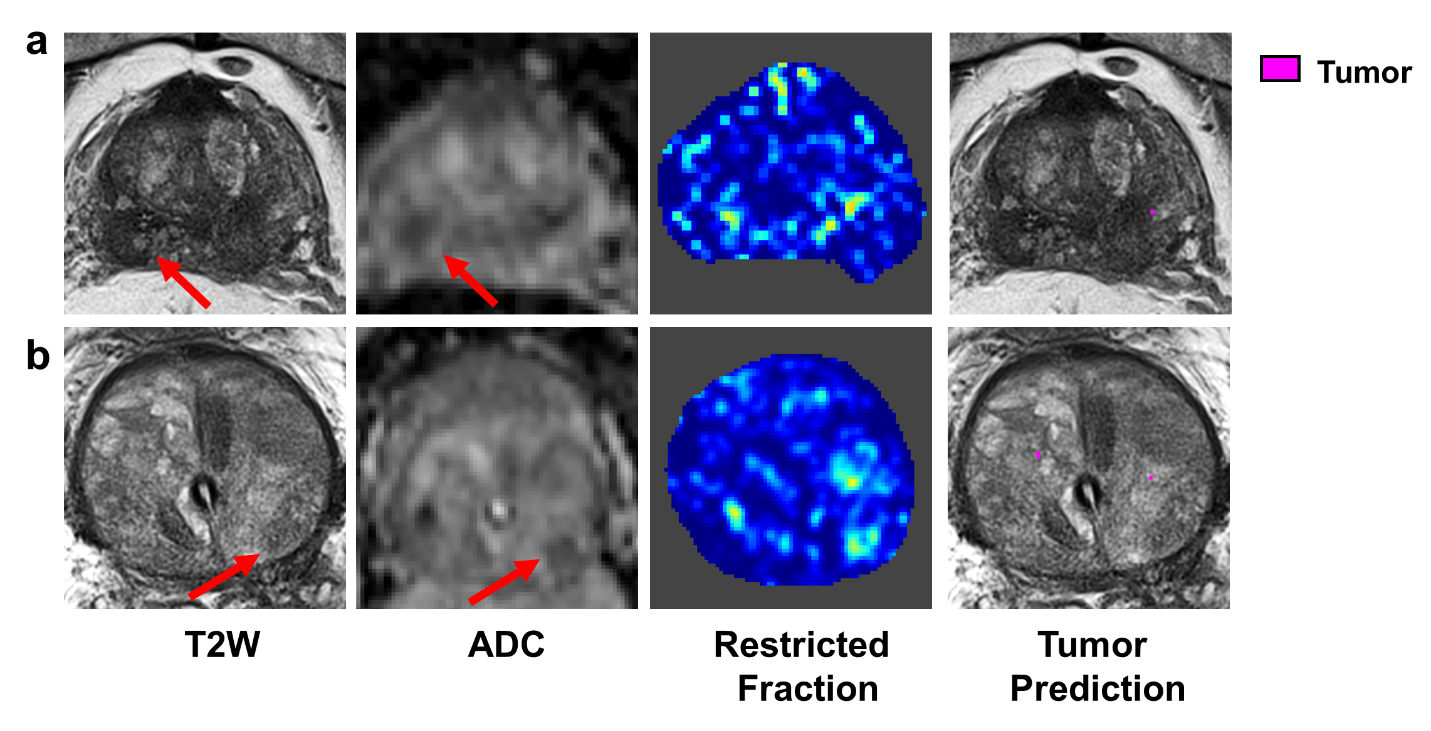

**Figure S3.** **False positive cases from mpMRI were corrrectly predicted by DHI.** Two representative cases were both PI-RAIDS v2 score 4 with suspicious lesions identified in ADC maps (a & b, arrows). DHI predicted these two patients to be benign, in consistent with biopsy results.
